# From raw reads to trees: Whole genome SNP phylogenetics across the tree of life

**DOI:** 10.1101/032250

**Authors:** Sanaa A. Ahmed, Chien-Chi Lo, Po-E Li, Karen W. Davenport, Patrick S. G. Chain

## Abstract

Next-generation sequencing is increasingly being used to examine closely related organisms. However, while genome-wide single nucleotide polymorphisms (SNPs) provide an excellent resource for phylogenetic reconstruction, to date evolutionary analyses have been performed using different ad hoc methods that are not often widely applicable across different projects. To facilitate the construction of robust phylogenies, we have developed a method for genome-wide identification/characterization of SNPs from sequencing reads and genome assemblies. Our phylogenetic and molecular evolutionary (PhaME) analysis software is unique in its ability to take reads and draft/complete genome(s) as input, derive core genome alignments, identify SNPs, construct phylogenies and perform evolutionary analyses. Several examples using genomes and read datasets for bacterial, eukaryotic and viral linages demonstrate the broad and robust functionality of PhaME. Furthermore, the ability to incorporate raw metagenomic reads from clinical samples with suspected infectious agents shows promise for the rapid phylogenetic characterization of pathogens within complex samples.

## INTRODUCTION

The reconstruction of evolutionary history using phylogenetics has been a fundamental method applied to many areas of biology. Single nucleotide polymorphisms (SNPs), one of the dominant forms of evolutionary change, have become an indispensable tool for phylogenetic analyses. Phylogenies in the pre-genomic era relied on SNPs found within single genes as the evolutionary signal, and later incorporated multiple loci, such as in multiple locus sequence typing (MLST). Although still valuable, these methods are limited to differences found in short sequences representing only a fraction of the genome, and are unable to capture the complete variation within species. As a result, these methods generally provide a low or an insufficient phylogenetic signal as gene-based trees do not always reflect the true species tree(Pamilo and Nei 1988). While phylogenetic analyses that use many orthologs are an improvement, multi-ortholog methods only utilize variable sites within annotated coding regions that pass a user-specified orthology test, and, more problematically, require both an accurate and existing annotation of all of the orthologous genes found within assembled genomes.

Whole-genome SNPs are one of the most commonly used features for measuring phylogenetic diversity since they can identify key phylogenetic clades for most organisms and can help resolve both short and long branches(Girault et al. 2014; Griffing et al. 2015). Additionally, since selectively neutral SNPs accumulate at a uniform rate, they can be used to measure divergence between species, as well as strains(Schork et al. 2000; Filliol et al. 2006). Furthermore, due to the large number of SNPs found along the length of entire genomes, the use of whole-genome SNPs minimizes the impact of sequencing and assembly errors, as well as individual genes under strong selective pressure.

Although genome-wide sequencing now allows examination of the entire genome, finished genomes are only rarely produced due to platform biases, or computational and cost limitations (e.g. 111,857 assembled genomes versus 7,555,153 SRA datasets in NCBI). However, comparative genomics, including SNP- and ortholog gene-based phylogenetic analyses usually require assembled or finished genomes(Ben-Ari et al. 2005; Foster et al. 2009). Here we present PhaME, a whole-genome SNP-based approach that can make use of available completed genomes, draft assemblies (contigs), and raw reads to perform <u>P</u>hylogenetic and <u>M</u>olecular <u>E</u>volutionary analyses.

Several methods for whole-genome SNP discovery or phylogenetics have been described: SNPsFinder(Song et al. 2005), PhyloSNP(Faison et al. 2014), kSNP(Gardner and Hall 2013), WG-FAST(Sahl et al. 2015), and CFSAN(Davis et al. 2015). SNPsFinder requires assembled genomes, uses the time-consuming megaBLAST program, and only provides a table of identified SNPs. PhyloSNP requires pre-aligned data (such as vcf files or tab-delimited lists of SNPs), does not scale to large datasets, and only produces maximum parsimony trees based on the presence or absence of SNPs. Like PhaME, kSNP, WG-FAST, and CFSAN can analyze raw reads to identify a core genome (the conserved portion among all genomes). However kSNP is restricted to finding central SNPs within an optimal kmer window, the size of which must first be provided by the user and influences the resulting tree. WG-FAST is a rapid method that identifies the most closely related known genome for given input data, but requires a pre-formatted SNP matrix of known SNP positions for a target organism and a pre-computed phylogeny. CFSAN aligns read datasets against a designated reference genome to generate a SNP matrix, but only allows the inclusion of a single reference genome and does not build phylogenies. Because each of these pipelines perform only part of the steps required to infer phylogenies from a particular form of sequencing data (i.e. either reads or contigs), we have developed an integrated PhaME analysis pipeline which provides rapid tree construction from assemblies and reads and downstream evolutionary analysis of functional genes, with the ability to incorporate pre-constructed alignments and phylogenies.

PhaME is an open source utility (https://github.com/LANL-Bioinformatics/PhaME) that combines algorithms for whole-genome alignments, read mapping, and phylogenetic construction, and uses in-house scripts to discover the core genome and core SNPs, infer trees, and conduct further molecular evolutionary analyses. Because of the algorithm’s design, PhaME can also provide phylogenetic analyses of a target organism present even at low abundance within metagenome samples, a feature unique to PhaME. This method is rapid and extensible, able to build upon an existing multiple sequence alignment for any set of related genomes and infer a new tree. Similarly, trees for subsets of the input genomes can readily be calculated. We demonstrate PhaME’s ability to construct robust phylogenies that include raw sequencing read datasets as input (including metagenome samples) by building phylogenies with up to 560 genomes and spanning the tree of life.

## RESULTS

### Implementation of PhaME across the tree of life

PhaME was designed to identify SNPs anywhere in the genomes of closely related organisms and to infer phylogenetic trees using any type of sequence input, including finished genomes, assembled contigs of draft genomes, or even FASTQ raw read datasets (see Methods). We tested PhaME on: 1) two large groups of bacteria, 2) the largest available dataset for a eukaryotic genus *(Saccharomyces)*, and 3) a viral dataset that includes many assemblies from the recent 2014 *Zaire ebolavirus* outbreak. We have further examined the robustness of how PhaME handles raw read datasets, including those from metagenomic samples that contain varied amounts of the target organism. Table 1 summarizes the number of genomes and raw reads used in the analyses, the average genome size per group, the calculated core genome sizes (and percent of the average genome size), the number of SNPs found in the core genome (and percent of the core genome size), and the number of coding (CDS) SNPs (and percent of the total SNPs), as well as the runtime information.

**Table 1.** Summary of genomes, assemblies and read datasets together with core genome statistics.

|  | *Escherichia/Shigella* | *Ecoli/Shigella* | Metagenome addition | *Escherichia/Shigella/Salmonella* | True *Escherichia/Shigella* | *Burkholderia* | *Bpseudomallei/Bmallei* | *Burkholderia BCC* | *Burkholderia BCC/pseudomallei/xenovorans* | *Saccharomyces* | *Scerevisiae* | Zaire ebolavirus |
| --- | --- | --- | --- | --- | --- | --- | --- | --- | --- | --- | --- | --- |
| Number of NCBI genomes (Complete and Draft) | 302 | 291 | 294 | 413 | 308 | 163 | 118 | 25 | 27 | 77 | 45 | 558 |
| Number of NGS read files | 0 | 0 | 1 | 0 | 0 | 31 | 49 | 0 | 0 | 2 | 0 | 2 |
| Total genomes | 302 | 291 | 295 | 413 | 308 | 194 | 167 | 25 | 27 | 79 | 45 | 560 |
| Average genome size | 4966643 | 4999028 | 4966643 | 4783138 | 4999028 | 6957651 | 6901660 | 7367266 | 7544548 | 12071326 | 12071326 | 18613 |
| Total gap length | 4904879 | 3331000 | 3620525 | 4986576 | 3686851 | 6279803 | 3956942 | 5478178 | 6071363 | 10929991 | 8262225 | 1398 |
| Total repeats | 180894 | 180894 | 180894 | 180894 | 180894 | 226210 | 226210 | 113966 | 113966 | 874683 | 874683 | 0 |
| Core genome size | 249983 | 1823862 | 1534337 | 168191 | 1468011 | 1028251 | 3351112 | 1800938 | 1207753 | 1141335 | 3809101 | 17561 |
| Total core SNPs | 73640 | 541834 | 371060 | 49337 | 461639 | 72968 | 89399 | 391765 | 274048 | 463 | 823064 | 1399 |
| CDS SNPs | 71341 | 502760 | 347746 | 47813 | 433890 | 72642 | 71479 | 380062 | 269380 | 463 | 664902 | 928 |
| % core of the genome | 5.03 | 36.48 | 30.89 | 3.52 | 29.37 | 14.78 | 48.56 | 24.45 | 16.01 | 9.45 | 31.55 | 94.35 |
| % total core SNPs of core | 29.46 | 29.71 | 24.18 | 29.33 | 31.45 | 7.10 | 2.67 | 21.75 | 22.69 | 0.04 | 21.61 | 7.97 |
| % CDS SNPs of total SNPs | 96.88 | 92.79 | 93.72 | 96.91 | 93.99 | 99.55 | 79.96 | 97.01 | 98.30 | 100.00 | 80.78 | 66.33 |
| Wallclock time |  |  | 0:02:06:28 | 1:02:39:06 |  | 0:14:13:44 |  |  |  | 0:15:32:36 | 0:1:26:48 |  |
| User time |  |  | 0:16:57:07 | 19:13:32:22 |  | 1:23:09:59 |  |  |  | 0:14:18:05 | 0:08:25:48 |  |
| System time |  |  | 0:01:40:06 | 0:14:29:11 |  | 0:00:59:09 |  |  |  | 15:17:0:35 | 0:01:09:19 |  |
| Maximum vmem |  |  | 9.892G | 50.720G |  | 66.394G |  |  |  | 51.405G | 8.479G |  |

#### Fine-scale phylogenetic analysis among closely related bacterial genera

The model bacterium *Escherichia coli* has been extensively studied, including its diversity and phylogenetic history. In previous studies, phylogenetic analyses using either 31 essential genes(Sahl et al. 2012) or all identified core genes(Ahmed et al. 2012) have shown that 1) the named *E. coli* species can be clustered into five main phylogenetic groups (A, B1, B2, D, E), 2) *Shigella sonnei, Shigella boydii*, and *Shigella flexneri* form two phylogenetic groups (groups SF and SS) within the *E. coli* phylogeny, and 3) *Shigella dysenteriae* clusters with the *E. coli* phylogenetic group E. These analyses demonstrate that the *Shigella* species are closely related to *E. coli* and not truly a separate genus(Chaudhuri and Henderson 2012), and that other *Escherichia* species (E.g. *E. blattae)* are likely inaccurately classified as *Escherichia*(Priest and Barker 2010). More recently, environmental organisms assigned to *Escherichia* appear to form independent lineages, or ‘cryptic clades’ and are also closely related to *Salmonella*(Walk et al. 2009). While these studies have illustrated the diversity of *Escherichia*, each study has imposed a different method of phylogenetic analysis sometimes resulting in incongruent trees(Walk et al. 2009; Luo et al. 2011), and none of the studies incorporates strains of all lineages or are able to provide a high-resolution phylogeny. We used this well-characterized group to both validate PhaME and to help better understand the phylogenetic relationships among members of this group at a more granular level than previous studies.

Using a total of 413 NCBI complete and draft enteric genomes (including *Shigella, Escherichia* and *Salmonella*; Supplemental Table S1), PhaME calculated a core genome of 168,191 bp with 49,337 SNP positions (of which 47,813 resided within annotated coding sequences; Table 1) and inferred a phylogenetic tree (Figure 2; Supplemental Fig. S1). Despite the inclusion of distantly related organisms, the phylogeny shows an identical topology for the *E. coli* and *Shigella* clades compared with prior studies(Ahmed et al. 2012; Sahl et al. 2012). Additionally, this tree resolves the evolutionary relationships among the environmental cryptic *Escherichia* lineages. For example, consistent with the 2009 MLST study, but in contrast with the 2011 single copy core gene study, the *E. albertii* lineage diverged before *E. fergusonii*(Walk et al. 2009; Priest and Barker 2010). In addition, this tree also supports the renaming of *E. blattae* to *Shimwellia blattae*(Walk et al. 2009), and further suggests that *E. hermanii* should also be reclassified into a different genus.

**Figure 1.**
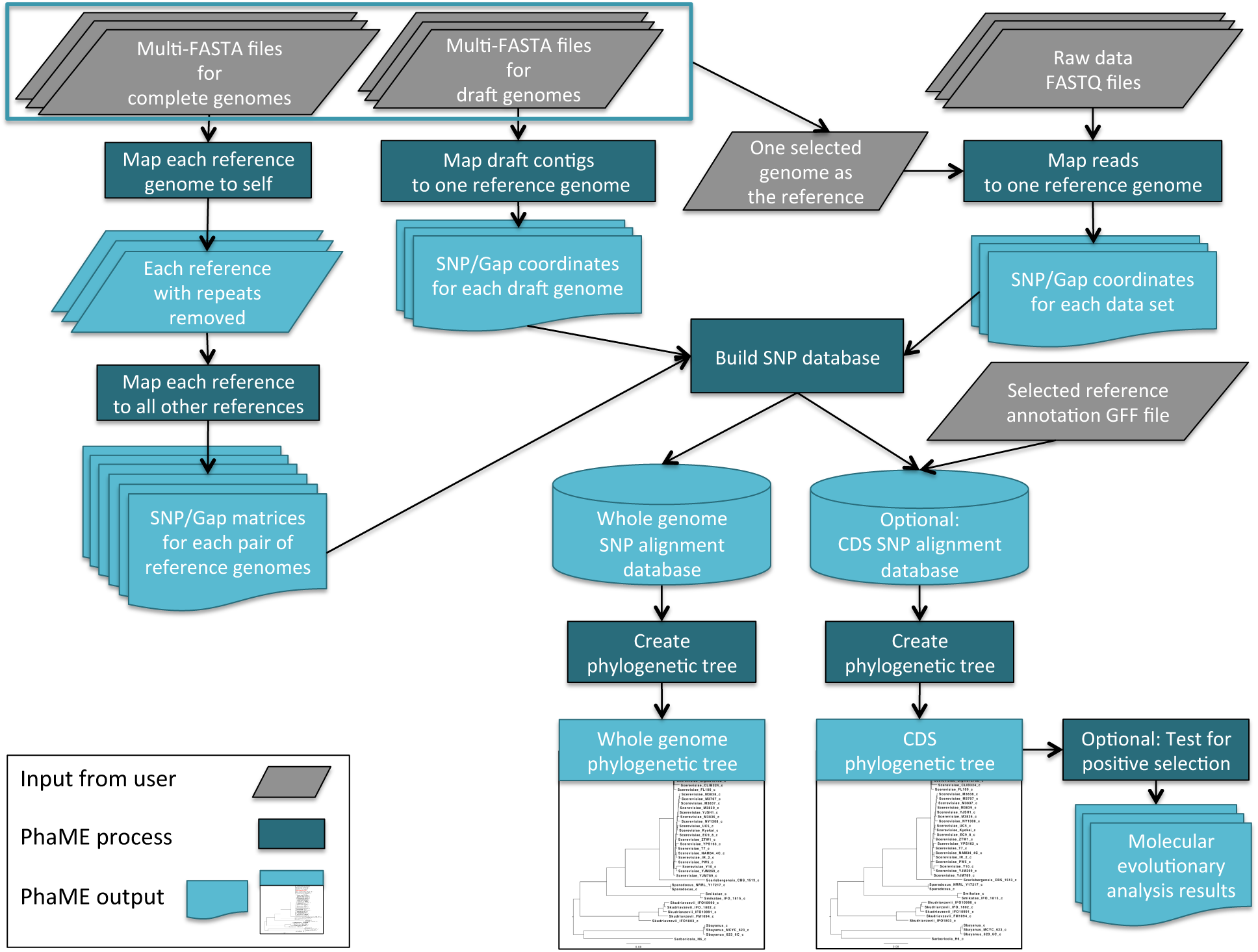
PhaME analysis pipeline. The PhaME analysis pipeline is able to identify SNPs from complete, assembled, and read datasets and infer a phylogenetic tree. If assembled genomes are provided, NUCmer is used to identify repeats and perform pairwise alignments. Bowtie2 or BWA is used to map reads to one of the repeat-masked genomes. The SNP and gap coordinates are used to generate whole-genome SNP alignments. If an annotation file is provided, a separate alignment consisting of SNPs only found in the CDS regions is generated. RAxML or FastTree phylogenies are constructed using the SNP alignments. If specified, PAML or HyPhy packages can be used to test for episodic diversifying selection.

**Figure 2.**
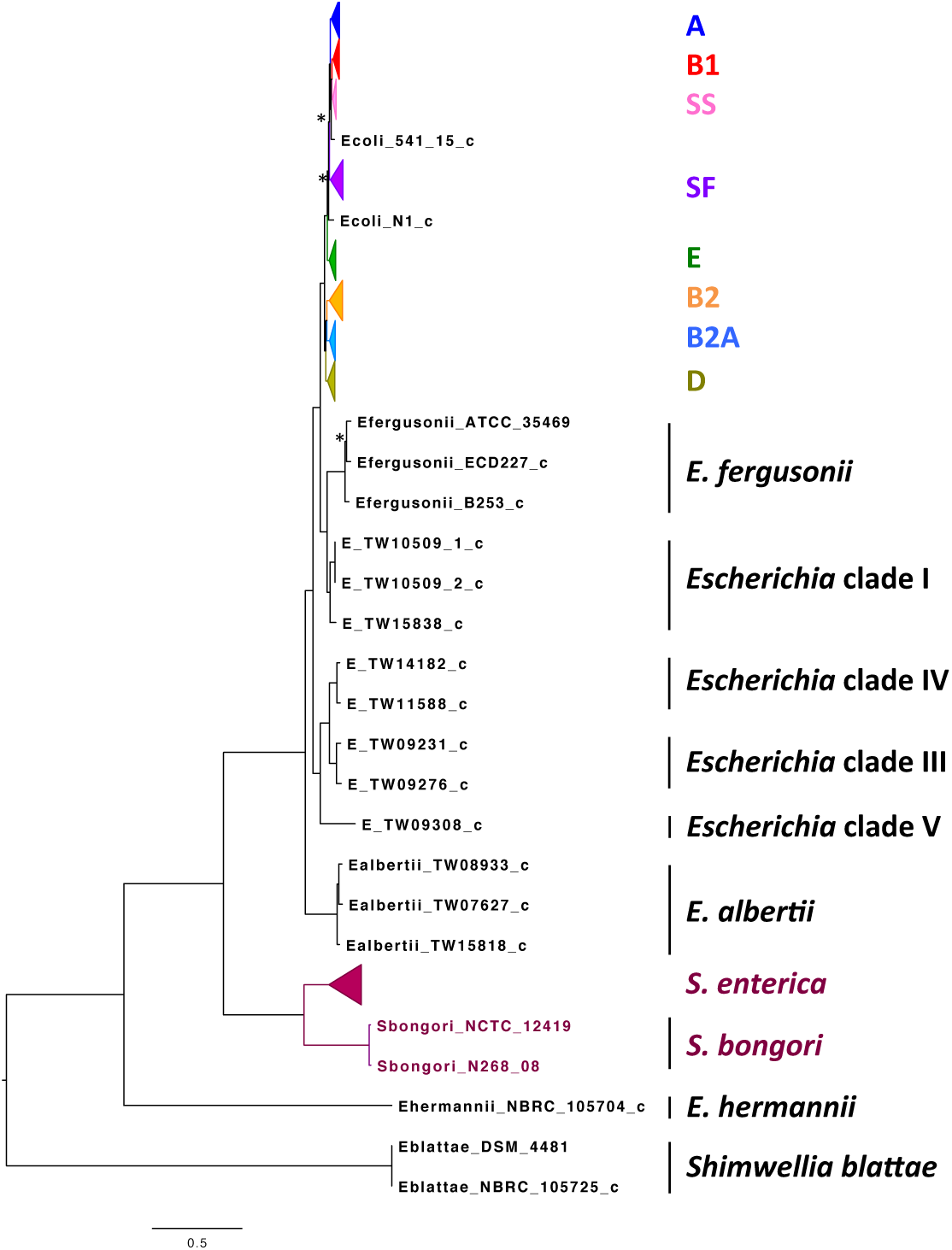
Inter-genus phylogeny of *Escherichia, Shigella* and *Salmonella* strains. Whole-genome SNPs from 413 *Escherichia*, *Shigella*, and *Salmonella* genomes were used to construct a maximum likelihood phylogenetic tree. *Shigella* and *E. coli* strains cluster together into previously identified phylogroups (colored, collapsed clades), with the *Salmonella* genomes as a distinct outgroup. Stars (*) represent bootstrap values under 75, all other clades are supported by values >75. Entries with _c represent contig datasets, the remaining datasets are finished genomes. See Supplemental Figure S1 for a fully detailed phylogenetic tree of this group. Scale = 0.5 substitutions per base.

While this inter-genus tree helped determine the relationships among these divergent lineages, the available SNP matrix (see Methods) allows PhaME to rapidly perform a more refined analysis of any genome subset, for example, just the *Escherichia* genus (Supplemental Fig. S2), or just the *E. coli* clade (Supplemental Fig. S3). The core genome for all *Escherichia* (excluding the renamed *S. blattae* and *E. hermanii)* was calculated to be 1,468,011 bp, with 461,639 SNP positions, roughly ten times larger than the core (and core SNPs) found when including the other genera. The core genome for all *E. coli* (and *Shigella)* genomes is 1,823,862 bp with 541,834 SNP positions, providing even more resolution within those lineages. While the improved resolution of subtrees likely provides more accurate estimates of lineage evolution (branch lengths), no major topological differences were observed when comparing the original tree with the subtrees, supporting the robustness of PhaME even across genera.

#### Establishing robust placement of read datasets within a phylogeny

The *Burkholderia* genus is a highly diverse group, occupying a wide range of ecological niches, and can be either free-living or symbionts. Some members are commensals or pathogens of a variety of eukaryotic organisms. On a genome level, *Burkholderia* species have either 2 or 3 designated chromosomes whose sizes can vary tremendously. While the smallest genome, for *B. rhixozinica*, has 1 chromosome and 1 megaplasmid with a total genome size of 3.6 Mbp(Lackner et al. 2011), *B. xenovorans* has a much larger genome of 9.8 Mbp spanning 2 chromosomes and a megaplasmid(Daligault et al. 2014). As of 2009, there are over 30 described species of *Burkholderia*(Coenye and Vandamme 2003), many of which have a sequenced representative, (the biothreat pathogens *B. mallei* and *B. pseudomallei* are overrepresented among the sequenced genomes). One group commonly known as the *B. cepacia* complex (*Bc*c) was first described as a cluster of 9 closely related species and now contains 15 distinct species. Over the years, the *Bc*c have been uncharacteristically difficult to discriminate using conventional polyphasic, 16S, or MLST approaches, and there are still longstanding issues in resolving their evolutionary history(Coenye and Vandamme 2003; Storms et al. 2004; Reik et al. 2005; Baldwin et al. 2007; Vanlaere et al. 2009).

We used PhaME to infer a genus-level phylogenetic tree (Figure 3; Supplemental Fig. S4) using 163 complete or draft genomes as well as 31 read files (totaling 194 datasets and 0.783 TB of information; Supplemental Table S2). PhaME calculated a core genome for the *Burkholderia* genus of 1,028,251 bp with 72,968 total SNPs (of which 72,642 are found in coding sequences). In comparison with the *Escherichia* genus (excluding *Shimwella blattae* and *E. hermanii)*, this core is smaller with respect to the average genome size and the number of SNPs is proportionally even smaller. This confirms that the *Burkholderia* are highly diverse with a large accessory genome(Chain et al. 2006), but that the smaller core genome itself is highly conserved.

**Figure 3.**
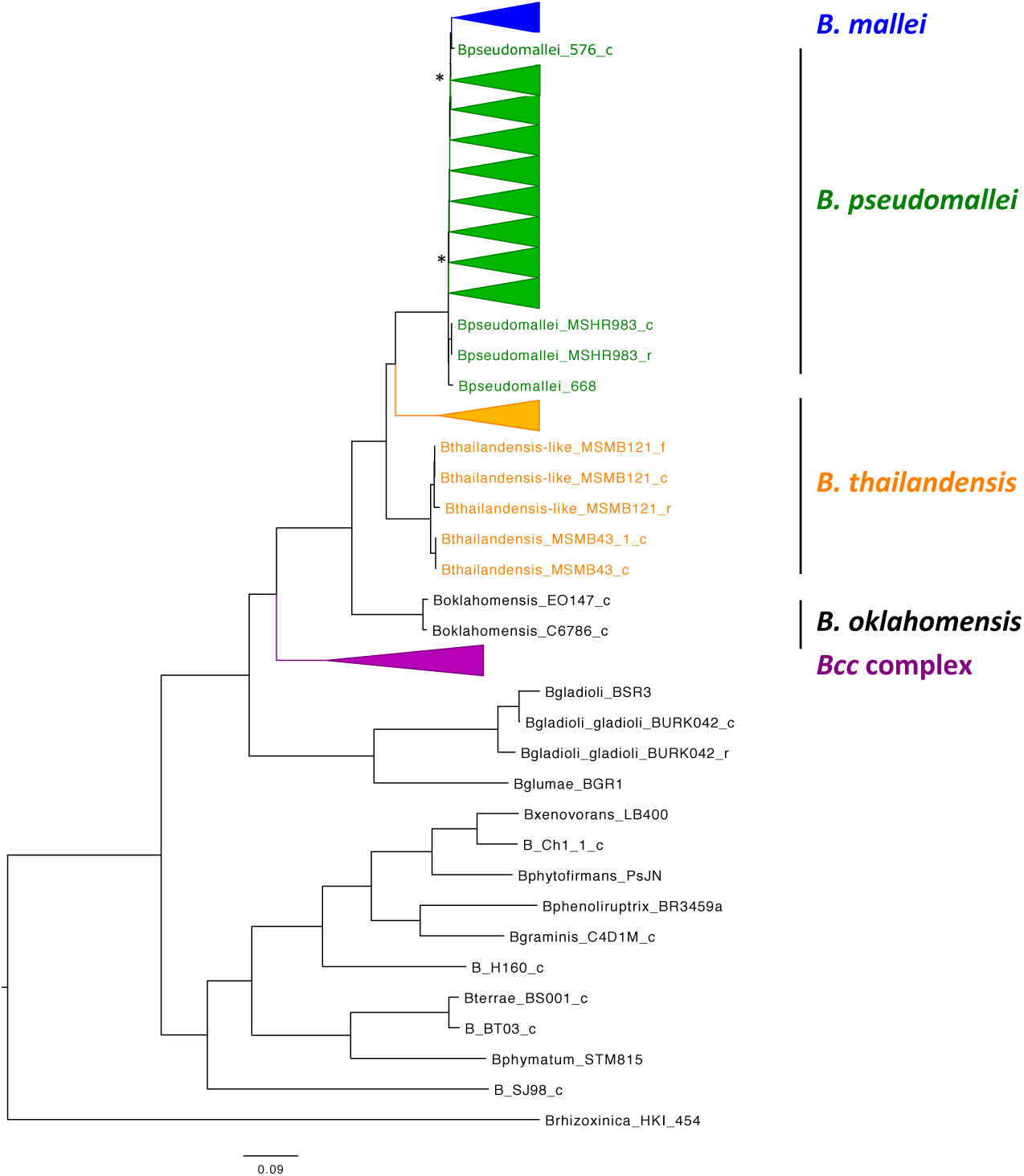
Phylogenetic tree of the *Burkholderia* genus using reads, contigs and finished genomes. Maximum likelihood phylogeny of 217 *Burholderia* genome datasets using PhaME. Collapsed branches represent various clades of the mentioned groups. Stars (*) represent bootstrap values under 75, all other clades are supported by values >75. Supplemental Figure S4 displays the detailed phylogenetic tree with all 217 entries, and demonstrates the placement of full genomes, assemblies and reads, which generally cluster together. Supplemental Figure S5 shows the relationships among the *Bcc* genomes together with outgroups. Supplemental Figure S7 provides a more detailed view of the placement of reads with contigs and genomes within the *B. pseudomallei* and *B. mallei* clade. Entries with _f, _c, _r represent: _f, finished genomes; _c, contigs from assembled genomes; r, raw reads. Scale = 0.09 substitutions per base.

The phylogenetic tree (Figure 3; Supplemental Fig. S4) highlights the recent clonal derivation of *B. mallei* from *B. pseudomallei*, with *B. pseudomallei* 576 as the most closely related genome, and recapitulates the paraphyletic nature of the *B. pseudomallei* strains(Godoy et al. 2003) when *B. mallei* is considered its own species. This tree also shows two well-supported (bootstrap value of 99) separate clades representing the *B. thailandensis* genomes suggesting a possible reclassification of one of these two groups as a separate species. In addition, while *B. gladioli* has been described as closely related to the *Bcc* species(Brisse et al. 2000; Coenye et al. 2001), not only is it monophyletic with *B. glumae*, but together these two species are found as an outgroup to the rest of the *Bc*c genomes and the *B. pseudomallei* group (further supported by the subtree in Supplemental Fig. S5). The placement of the endosymbiont *B. rhizoxinica* as an outgroup with respect to the environmental isolates was determined with PhaME when including three genomes of *Rastonia pickettii* as outgroup (Supplemental Fig. S6).

The ability to select a subset of genomes for analysis can provide insight into not only the consistency of the topology with the original tree, but can show differences in the resulting core size and the SNPs within the core. For example, while the topology of the *Bc*c complex subtrees remains the same as the original tree, the core genome size increases to 1,800,938 bp with 391,765 SNPs (380,062 within CDS). However, when examining only the *B. pseudomallei* (including *B. mallei)* lineage, the core genome is even larger (3,351,112 bp) yet the number of core SNPs is much smaller (89,399 with only 71,479 within CDS). This is in sharp contrast with the *Escherichia* and the *E. coli* core genomes and SNP counts, where an increase in core genome size is accompanied by an equivalent increase in SNPs. Furthermore, the number of SNPs used to infer the *Burkholderia* genus tree and the *B. pseudomallei* tree are similar, yet the *B. pseudomallei* tree provides much more discriminatory power among these closely related strains. Given these patterns, we reason that the nucleotides at SNP positions used for the *Burkholderia* genus tree are in fact identical among the *B. pseudomallei* genomes, and are not the same as those used for *B. pseudomallei* tree.

Because one of the other unique features of PhaME, beyond rapid subtree inference, is that it allows the inclusion of raw read datasets into a whole genome SNP phylogeny, we evaluated the accuracy of placement of read datasets compared with the resulting assemblies and finished genomes. Both the whole genus tree (Supplemental Fig. S4) and a more detailed *B. pseudomallei/B. mallei* subtree (Supplemental Fig. S7) correctly place the reads with the contigs and genomes that were derived from those reads (phylogenies using only the reads and not the contigs/genomes resulted in consistent topologies, data not shown). This self-consistency supports the ability of PhaME to conduct robust phylogenetic analysis without the need for assembling raw sequencing data prior to inclusion in a tree.

#### Application of PhaME to eukaryotic genomes

Because PhaME should be readily applied to any other group of closely related genomes, we decided to implement tests beyond bacterial lineages. Fungi are known to be a difficult group in terms of phylogenetic analysis(James et al. 2006a; James et al. 2006b; Hibbett et al. 2007). The phylogenetic placement of fungal species display disparities between trees based on gene sequence analyses and those based on morphological characteristics (such as modes of reproduction). This is especially true of the ‘*Saccharomyces* complex’, where 18S and 26S rDNA comparisons do not show well-supported clades, resolving only the most closely related species(Kurtzman and Robnett 2003).

Due to the complexity of assembling eukaryotic genomes, there are few reference draft or complete genome assemblies for eukaryotic species. This is the case for the *Saccharomyces* genus, which only has a single complete reference genome. For eukaryotic genomes, the ability to make use of raw reads can be of great value in characterizing those genomes. A total of 79 genome projects consisting of 1 complete genome for *S. cerevisiae S288c* (16 chromosomes that total 12,071,326 bp in length), 76 draft genomes, and 2 raw read sets (Supplemental Table S3) were used as input to the PhaME pipeline. All 16 chromosomes of the yeast genome were used for this analysis and, using all the input datasets, we generated the first whole genome phylogeny using all available *Saccharomyces* genomes. This phylogenetic tree is based on only 463 whole-genome SNP positions (explaining some of the low bootstrap confidence values in the tree), all of which reside within CDS regions identified in 1,141,335 bp of core genome (Supplemental Fig. S8). This also highlights the highly conserved nature of a eukaryote’s core genome. A refined analysis focusing solely on the *S. cerevisiae* clade increases the core genome size to 3,809,101 bp, consisting of 823,064 whole-genome SNPs, with 664,902 CDS SNPs. With such a dramatic increase in the number of positions used for tree inference, one can observe much improved discrimination among strains, with stronger bootstrap support for most ancestral nodes (Supplemental Fig. S9). This analysis is a novel approach for eukaryotic genomes, where whole genome SNPs were used for phylogenetic inference instead of solely relying on gene annotations. This method allows for the robust discrimination among *S. cerevisiae* strains and helps better understand the relationships among *Saccharomyces* species.

#### Whole genome PhaME analysis of a viral outbreak

The *Zaire ebolavirus* outbreak that began in 2014 was rapidly characterized by large-scale sequencing and assembly of *Zaire ebolavirus* genomes from several hundred patients(Gire et al. 2014; Carroll et al. 2015; Simon-Loriere et al. 2015). Because PhaME can provide exquisite resolution among members of the same species, it could be useful in viral pathogen outbreak investigations. Therefore, PhaME was applied to a total of 538 assembled genomes sequenced from the 2014/2015 *Zaire ebolavirus* outbreak, as well as 20 reference genomes sequenced from previous outbreaks (Supplemental Table S4). PhaME calculated 17,561 bp as the core genome size (which approaches the average *Zaire ebolavirus* genome size of ~18,959 bp) with 1399 core SNP positions, of which 928 are found in annotated coding regions. The PhaME *Zaire ebolavirus* phylogeny clearly distinguishes the previous outbreaks from the 2014 outbreak (Supplemental Fig. S10). Additionally, this topology is well supported and consistent with the previous trees proposed from this outbreak(Gire et al. 2014; Carroll et al. 2015; Simon-Loriere et al. 2015). As previously observed, the *Zaire ebolavirus* strains recovered from Guinea and Sierra Leone are interspersed throughout the tree yet form some distinct lineages(Carroll et al. 2015; Simon-Loriere et al. 2015).

#### Analyzing raw metagenomic reads with PhaME

Outbreaks such as the case of *Zaire ebolavirus* provide a scenario where assembly of genomes is often the first step in analysis due to the fact that obtaining a pure isolate may be difficult or take longer than desired for rapid analysis. Since PhaME can accurately identify where raw read datasets belong in a phylogenetic tree (as shown above for reads sequenced from pure cultures), we tested PhaME’s ability to accurately place a known infectious agent within a phylogeny, using reads derived from complex metagenomic samples. We hypothesize that a dominant clonal pathogen infecting a host will be accurately placed within a phylogeny due to the read mapping and SNP calling strategy in PhaME. To our knowledge, this is the first demonstrated use of raw shotgun metagenomic reads (or any metagenomic data) as an input to phylogenetic reconstruction software. We investigated samples taken from individuals afflicted during the 2014 *Zaire ebolavirus* outbreak, as well as a clinical fecal sample from a US patient having returned from Germany during the 2011 stx2-positive Enteroaggregative *E. coli* (StxEAggEC) outbreak.

For *Zaire ebolavirus* metagenomes, we selected two 2014 datasets from Sierra Leone (both have few to no reads that can be matched to human sequences), one (SRX674125) with 3,505,216 reads, of which ~62.2% map to the *Zaire ebolavirus Mayinga 1976* reference sequence (2,179,715 reads; 8,861.13 average fold coverage), and another (SRX674271) with 929,604 reads, of which only 0.3% can be mapped to the same reference (2,850 reads; 9.45 average fold coverage). In both cases, assembled genomes derived from these two datasets are available in Genbank (Supplemental Table S4) and were also used in the PhaME *Zaire ebolavirus* tree (Supplemental Fig. S10). Adding the two full raw metagenome datasets to the *Zaire ebolavirus* tree took 1 hour 21 minutes, did not impact the core genome or SNP statistics (Table 1), and were both placed within the 2014 outbreak lineage. The sample with high *Zaire ebolavirus* load was positioned very closely to the sample-matched assembled genome (Figure 4, Supplemental Fig. 11). The other sample was placed as the immediate outgroup to all other 2014 strains, likely stemming from the paucity of data within this dataset. However, these results highlight the power of PhaME to phylogenetically characterize a target organism that comprises only a minute fraction of a complex sample.

**Figure 4.**
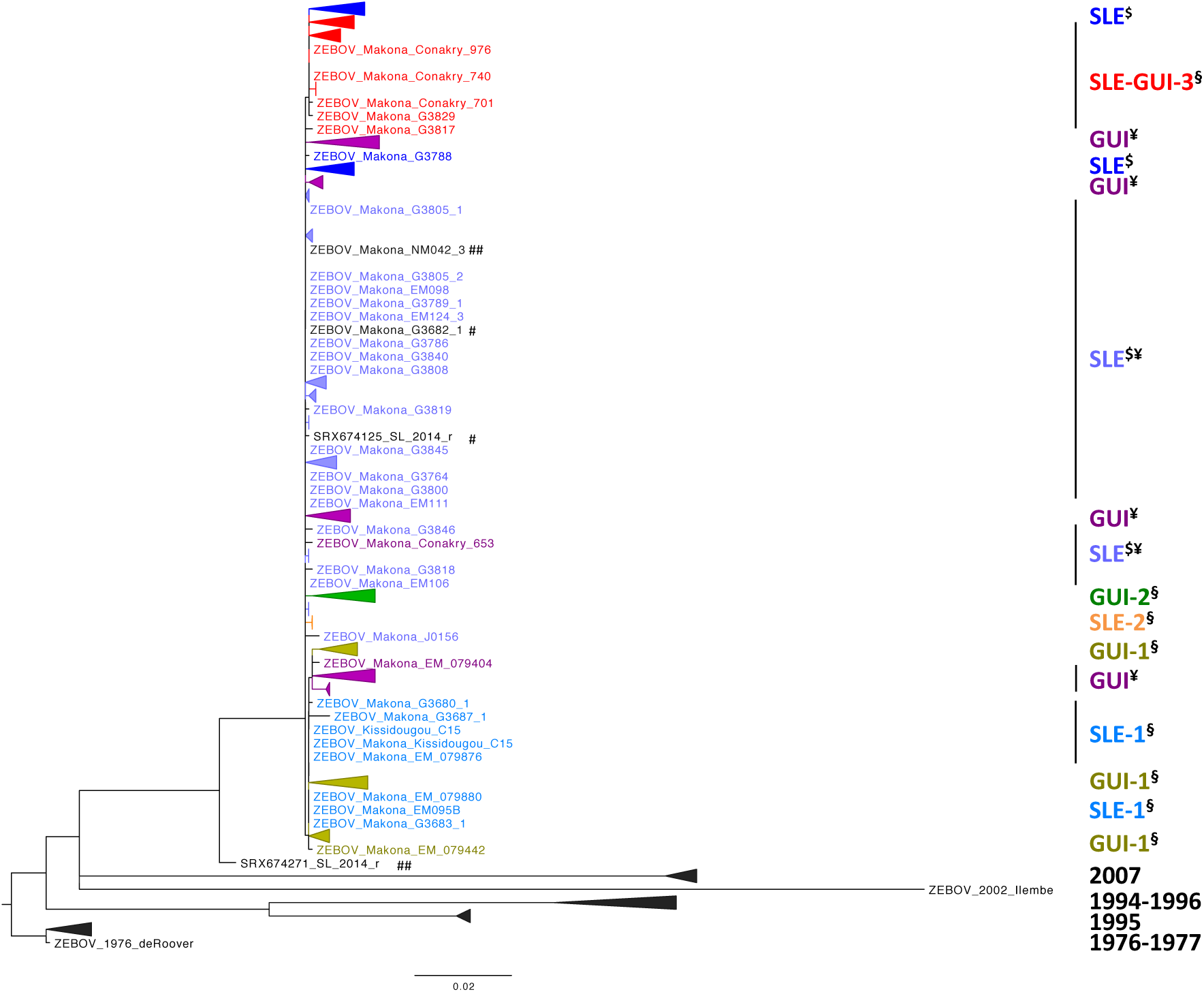
Accurate placement of metagenomic samples within the *Zaire ebolavirus* phylogeny. PhaME was used to generate a maximum likelihood phylogeny of 560 *Zaire ebolavirus* genomes (20 reference genomes, 538 assembled genomes from 2014 *Zaire ebolavirus* outbreak, and two raw metagenomic read datasets from individuals suspected of having been infected in this outbreak). Collapsed branches represent various clades of *Zaire ebolavirus* that are consistent with those reported previously(Gire et al. 2014)(Carroll et al. 2015). Supplemental Figure S11 displays the detailed phylogenetic tree with all 560 entries. Entries with _r represent raw metagenomic read datasets. Entries labeled as: ^$^ represent data from reference(Gire et al. 2014); ^¥^ represent data from reference(Carroll et al. 2015); ^§^ represent data from reference(Simon-Loriere et al. 2015). Scale = 0.02 substitutions per base.

In the context of metagenomic data, the ability to accurately phylogenetically place a target genome requires that a dominant clonal member of the target lineage be present in the sample in order to accurately identify SNPs. With *E. coli* as a commensal resident within the human gut, we tested the ability of PhaME to analyze a fecal sample derived from a patient suspected to be infected with the 2011 StxEAggEC strain. A large (252,926,569 reads) fecal sample dataset was mapped to 52 *E. coli* strains to evaluate the distribution of reads among available genomes. While all genomes recruited reads from this dataset (total of 1.74 million reads), the dominant organism within the sample appeared to belong to the StxEAggEC outbreak clade (Supplemental Fig. S12). Using the existing *E. coli+Shigella* PhaME phylogeny, the addition of the fecal sample’s raw shotgun metagenomic reads reduced the *E. coli* core genome size, as well as the core SNPs (Table 1). The dominant *E. coli* within the fecal sample was accurately placed in the inferred tree within the StxEAggEC phylogroup B1 (Supplemental Fig. S13), further supporting the use of PhaME to establish the phylogenetic placement of infectious disease outbreak strains, even when in the presence of less abundant commensal strains of the same species.

### Molecular analyses and signs of positive, negative and neutral selection

Identifying SNPs found in coding regions enables further molecular evolutionary analyses -a post-phylogeny option that is provided in PhaME. PhaME uses the HyPhy package Branch-Site random effects likelihood (REL) model for detecting episodic diversifying selection on all genes containing at least 1 SNP, as well as on a concatenation of all these genes.

The *Escherichia* and *Shigella* core genome was used to illustrate the application of molecular evolutionary analyses with PhaME. A total of 324 genes were found to contain at least one SNP, of which only 52 genes were detected to have lineages that showed a statistically significant signal of diversifying selection (Supplemental Table S5). Only four genes showed signs of positive selection, three of which were only found in the avian pathogenic *E. coli* within the tree. Because the size of the core genome decreases with the addition of more genomes, the core genome becomes increasingly enriched in essential genes and depleted in accessory genes. The small size of the core (~5% of the average genome size) and the high number of essential genes within the core genome may partially explain why so few genes show signs of diversifying selection. We examined the ability of PhaME to explore a single lineage within the larger phylogeny, consisting of a subset clade of 20 *Escherichia* and *Shigella* genomes. In this case, a total of 2513 genes were found to contain at least one SNP and, in 111 instances, a gene showed a statistically significant signal of diversifying selection, either in a single strain or within a monophyletic lineage comprising multiple strains. While nine genes of varied function showed signs of positive selection (Supplemental Fig. S14), most genes are under negative selection, including a number of genes involved in metabolism and housekeeping functions (Supplemental Table S6).

## DISCUSSION

With the rapidly growing number of available genomes and NGS read datasets, it is becoming increasingly important to have appropriate analysis tools that can deal with both assembled contigs (or complete genomes), as well as raw sequencing data. Several considerations in developing tools include: 1) the ability to handle short reads with errors while still producing accurate results, and 2) the ability to process large amounts of data in reasonable timeframes. It is also becoming increasingly important that tools be designed modularly to accommodate different research goals and that the tools be widely applicable. Here, we describe a new phylogenetic tool, PhaME, that can rapidly process hundreds of genomes and/or raw reads to obtain highly robust, whole genome SNP phylogenetic trees, and to estimate molecular evolution along lineages. This tool is novel due to its ability to: 1) incorporate both raw read datasets (including metagenomic) and genome assemblies, 2) uniquely combine an automatic core genome search, SNP identification and phylogenetic tree generation, and 3) assess the selective pressure along lineages.

We have tested PhaME using viral, eukaryotic, and bacterial genomes, and have constructed trees that include many hundreds of genomes of the same genus, as well as different but related genera. Using these examples from across the tree of life, we have demonstrated the ability to rapidly process up to almost 1 TB of data and to construct highly robust phylogenies, some of which have never before been done at this scale. When given an annotation for one of the reference genomes, we have also shown the ability to perform molecular evolutionary analyses. Although inclusion of this option significantly increases the runtime and memory required, it can be used to assess positive selection in distinct lineages, and can lead to hypotheses based on biologically relevant genomic signals.

We have shown PhaME’s ability to construct phylogenies with the inclusion of raw metagenomic data, a feature unique to PhaME. In two examples using sequenced clinical samples from independent outbreaks, one with *Zaire ebolavirus* and one with *E. coli*, the placement of the sample within the context of the other strains use in the phylogeny support assumptions regarding the identity of the pathogen found within the samples. In the *Zaire ebolavirus* example, with as little as 0.3% of the sample reads belonging to the actual pathogen, the placement within the tree lies within the correct clade of the published outbreak genomes. With *E. coli*, despite having a dominant presence of the StxEAggEC within the sample, significant amounts of data could be best matched with other (commensal) *E. coli*, presumably originating from the patient’s normal microflora. In this case, the robust placement in the tree is due the dominance of the target strain and the algorithms used to construct the alignment and phylogeny. Given the growing trend in sequencing metagenomes from ill individuals, this particular application of PhaME may be highly useful in a clinical setting.

We have included within PhaME the ability to incrementally add samples to a previously constructed core alignment (and tree), which allows for rapid analysis of additional samples, and have similarly included the ability to rapidly recompute the core genome and derived phylogenies using subsets of the input genomes. While phylogenetic analysis has traditionally required annotated genes, here we provide a highly automated process for today’s genomic era, agnostic of the input sequencing data that enables the construction of whole genome core alignment, phylogenetic, and molecular evolutionary analyses within a single tool.

## METHODS

### Modular Design and PhaME Overview

We present a tool for Phylogenetic and Molecular Evolution Analyses (PhaME) that can take raw NGS reads or assembled contigs representing draft or complete genomes, align the core conserved sections among the genomes, identify all SNPs (both coding and non-coding), infer a phylogenetic tree, and perform evolutionary analyses, such as identifying signals of positive selection. PhaME is a Perl program that incorporates several other open source software, including the MUMmer package NUCmer(Kurtz et al. 2004), Bowtie2 2.1.0(Langmead and Salzberg 2012), BWA 0.7.5a(Li and Durbin 2009), SAMtools 0.1.18(Li et al. 2009), HyPhy(Pond et al. 2005), PAML(Yang 2007), jModelTest(Darriba et al. 2012), RAxML(Stamatakis 2006), FastTree(Price et al. 2010), R package APE(Popescu et al. 2012). The overarching architecture of the PhaME analysis pipeline is outlined in Figure 1. PhaME is run using a control file (SNPphy.ctl) that can be modified by the user, and requires a minimum of one reference genome in FASTA format, consisting of one or more contigs (which can also be complete chromosomes, megaplasmids, plasmids, etc.). Additional genomes in the following formats can also be included in the analyses: finished genomes and draft contigs as FASTA files, and raw next generation sequencing reads in FASTQ format. The directory structure and output files created by the PhaME analysis pipeline are outlined in Supplemental Figure S15.

The main outputs of PhaME include all pairwise contig alignments, the core genome alignment (all orthologous positions conserved among all input genomes), the final alignment of all positions with one or more SNPs (a subset of the core), maximum likelihood tree(s), text files summarizing the number of SNPs in pairwise comparisons, the positions of all SNPs in all input genomes, and information on whether these SNPs alter a codon and its associated amino acid (for a full list of output files, see Supplemental Fig. S15). The molecular evolution analyses are performed on each gene that contains a SNP (when a GFF annotation file is provided for the reference) and are presented in a series of files per gene as well as in a summary file for all genes. Additional information on PhaME software can be found at https://github.com/LANL-Bioinformatics/PhaME.

### Whole genome alignments and SNP discovery from genomes, contigs and reads

Any input contigs (draft or complete genomes) are initially subjected to self-comparisons using NUCmer in order to remove duplicated regions or other highly similar ‘repetitive’ elements (length ≥ 100; identity 95) that could give misleading alignment results. Whole and draft genome sequences then undergo pairwise whole genome alignment (in all combinations) using NUCmer. Gap regions, consisting of unaligned segments ≥1 nucleotide are considered evolutionarily uninformative and are removed. Input raw read datasets (either single or paired-end) are then individually aligned to a selected reference using BowTie2 (default) or BWA. The alignment results are parsed using SAMtools and perl scripts to group identified SNPs by shared genomic location within the selected reference. An orthologous SNP alignment is created for each genome, contig, and/or read set, and contains the nucleotides at all positions that are found in all genomes, and where at least 1 genome differs at that position. Given an annotation file in GFF format, the pipeline can also distinguish SNPs present within coding sequence (CDS) regions from those present in intergenic spaces. The CDS SNPs are used to build a separate phylogenetic tree and will also be used for downstream evolutionary analyses (when selected). The SNPs identified in all pairwise genome alignments, as well as those identified using reads mapped to one of the genomes are available as text files. In addition, pairwise SNP profiles for i) the core genome, as well as for ii) the core coding genome and iii) the core intergenic genome (when annotation is provided) are also available. These SNP matrices allow for rapid recalculation of the core sequences for any subset of genomes, and then reconstruction of subtrees.

The user is provided a proximity cut-off option that excludes SNPs that are located within the proximity. This feature may prove useful if horizontal transfer has mobilized multiple SNPs from one genome to another, confusing the evolutionary signal(Foster et al. 2009; Croucher et al. 2011). If the user does not select a reference genome for the above analyses, one will be selected randomly from among those with GFF files. If no GFF files are present, then a reference is chosen at random, and the optional CDS-specific modules (including the molecular evolutionary analyses) will not be conducted. Example output files for each of the steps are available from https://github.com/LANL-Bioinformatics/PhaME.

### Phylogenetic Reconstruction

The SNP alignment, consisting of all SNP positions present in any genome within the identified core genome, is used to construct a phylogenetic tree. If a GFF annotation file is provided, an additional tree can be generated from the subset of SNPs found only within coding sequences, or only within the intergenic spaces. The phylogenetic trees are inferred with using FastTree (default) and/or the RAxML-HPC maximum-likelihood method. When RAxML is selected, jModelTest is first run to determine the best substitution model to use when inferring the trees. Both RAxML and FastTree produce a newick file that can be viewed using a display tool such as FigTree (http://tree.bio.ed.ac.uk/software/figtree/), Dendroscope (http://ab.inf.uni-tuebingen.de/software/dendroscope/)(Huson and Scornavacca 2012), Archeaopteryx (https://sites.google.com/site/cmzmasek/home/software/archaeopteryx), EvolView(Zhang et al. 2012), iTOL (http://itol.embl.de)(Letunic and Bork 2011), and Phylodendron (http://iubio.bio.indiana.edu/soft/molbio/java/apps/trees/). For convenience, we have included a basic tree view within a PDF file using jphylo (http://sourceforge.net/projects/jphylo/).

### Molecular evolutionary analyses

When selected, the first step in molecular evolutionary analyses is to root the above generated tree using the APE package in R. Using the reference GFF file, all homologous genes containing SNPs are used to test for positive or negative selection through the implementation of methods within the HyPhy or PAML packages. Both packages can test for the presence of positively selected sites and lineages by allowing the *d_N_/d_S_* ratio (ω) to vary among sites and lineages. The branch-site REL model in the HyPhy package (hyphy.org) is used to detect instances of episodic diversifying and purifying selection. If PAML is selected, the M1a-M2a and M7-M8 nested models are implemented (molecularevolution.org/software/phylogenetics/paml). In this latter case, the likelihood ratio test between the null models (M1a and M8) and the alternative model (M2a and M7) at a significance cutoff of 5% provides information on how the genes are evolving. The results for each gene are then summarized in a table containing information on whether the gene is evolving under positive, neutral, or negative selection, along with p-values. Using HyPhy, a phylogenetic tree with evolutionary information on each branch is generated as a postscript file (for each gene as well as for the concatenation of all genes). With PAML, the newick tree file is modified to incorporate evolutionary information on any branch that is found to be under positive selection. We opted to provide PAML as an option, however recommend using HyPhy (set as default) for genome-scale projects due to its speed and concise output.

### Data Access

GenBank accession numbers for the sequencing data and genomes used in this study can be found in Supplemental Tables 1-4. The PhaME software together with documentation can be found at https://github.com/LANL-Bioinformatics/PhaME.

## Acknowledgements

This work was supported in part by the U.S. Defense Threat Reduction Agency’s Joint Science and Technology Office (DTRA J9-CB/JSTO) under contract number CB10152 to PSGC. We thank the LANL Genome Programs group, and in particular Jason Gans for reading and providing feedback with the manuscript. The authors report no conflict of interests.**Author contributions**: P.S.G.C. conceived the study. S.A.A. was responsible for algorithm design and bioinformatics analyses. C.L. was responsible for read mapping and alignment to references. P.L. and C.L. were responsible for making the pipeline available online. K.W.D. generated the flowchart for pipeline design. P.S.G.C. and S.A.A. interpreted the data and wrote the manuscript with input from the other authors.

